# Intraspecific trait changes have large impacts on community functional composition but do not affect ecosystem function

**DOI:** 10.1101/2021.02.05.429745

**Authors:** Noémie A. Pichon, Seraina L. Cappelli, Eric Allan

## Abstract

1. Plant functional traits can provide a more mechanistic understanding of community responses to global change and effects on ecosystem functions. In particular, nitrogen enrichment shifts trait composition by promoting dominance of fast growing, acquisitive plants (with high specific leaf area [SLA] and low leaf dry matter content [LDMC]), and such fast species have higher aboveground biomass production. Changes in mean trait values can be due to a shift in species identity, a shift in species relative abundance and/or a shift in intraspecific trait values. However, we do not know the relative importance of these three shifts in determining responses to global change and effects on function.
2. We quantified the relative importance of composition, abundance and intraspecific shifts in driving variation in SLA and LDMC. We collected leaf samples in a large grassland experiment, which factorially manipulates functional composition (slow vs. fast species), plant species richness, nitrogen enrichment and foliar fungal pathogen removal. We fitted structural equation models to test the relative importance of abundance shifts, intraspecific shifts and sown trait composition in contributing to overall variation in community weighted mean traits and aboveground and belowground biomass production.
3. We found that intraspecific shifts were as important as abundance shifts in determining community weighted mean traits, and even had large effects relative to a wide initial gradient in trait composition. Intraspecific trait shifts resulted in convergence towards intermediate SLA, in diverse communities, although convergence was reduced by nitrogen addition and enhanced by pathogen removal. In contrast, large intraspecific shifts in LDMC were not influenced by the treatments. Belowground biomass was reduced by SLA and increased by LDMC, while aboveground biomass increased in communities dominated by high SLA species. However, despite large intraspecific trait shifts, intraspecific variation in these traits had no effect on above or belowground biomass production.
4. Our results add to a growing body of literature showing large intraspecific trait variation and emphasise the importance of using field sampled data to determine community composition. However, they also show that intraspecific variation does not affect ecosystem functioning and therefore trait response-effect relationships may differ between vs. within species.

## Introduction

The use of functional traits has revolutionised community ecology and allowed predictions of how plant communities respond to environmental change and influence ecosystem functioning (McGill, Enquist, Weiher, & Westoby, 2006; Lavorel & Grigulis, 2012; de Vries et al., 2012; Allan et al., 2015). Understanding the correlations between trait responses to the environment and effects on functioning has been referred to as a “holy grail” (Lavorel & Garnier, 2002), which would allow a more predictive understanding of global change impacts. One of the key axes of plant functional trait variation is the leaf economics spectrum (Wright et al., 2004; Díaz et al., 2016), which distinguishes slow growing species with low leaf nutrients, tough and long-lived leaves from fast growing species with high leaf nutrient contents, soft and short-lived leaves. It is indicated by traits such as specific leaf area (SLA), leaf dry matter content (LDMC) and leaf nitrogen content, which tend to be strongly correlated with each other. These leaf economics spectrum traits predict responses to soil fertility, with increases in mean SLA and decreases in mean LDMC of communities in high nitrogen conditions (Lavorel & Grigulis, 2012; Allan et al., 2015). Leaf economics spectrum traits also predict effects on a number of ecosystem functions, as slow growing communities (with low SLA and high LDMC) typically produce less aboveground biomass and have slow biogeochemical cycling, but may allocate more resources belowground (see Freschet, Swart, & Cornelissen, 2015; Mahaut, Fort, Violle, & Freschet, 2019). Understanding the ways in which community trait values change in response to the environment and affect ecosystem functioning is critical to a mechanistic understanding of global change effects on plant communities and their functioning.

Community functional composition is typically measured as the mean trait value across all individuals, i.e. the community weighted mean, CWM, which is calculated by multiplying species trait values by their relative abundance. Shifts in community functional composition with environmental change can occur through three mechanisms: changes in species identity, a shift in the relative abundances of species with particular traits (hereafter abundance shift) or an intraspecific shift in traits between individuals (intraspecific shift) (Lepš, de Bello, Šmilauer, & Doležal, 2011). Intraspecific shifts can occur either through plastic changes in trait expression or through genetic changes (Geber & Griffen, 2003). It is commonly assumed that intraspecific shifts will have a smaller effect than abundance shifts on community mean trait values (Garnier, Laurent et al., 2001; Albert, Grassein, Schurr, Vieilledent, & Violle, 2011; Siefert et al., 2015), but, experiments have revealed large variation intraspecific variation in certain functional traits (Jung, Violle, Mondy, Hoffmann, & Muller, 2010; Violle et al., 2012; Albert, 2015), particularly specific leaf area (SLA) (Mitchell, Wright, & Ames, 2017; Derroire, Powers, Hulshof, Cárdenas Varela, & Healey, 2018). Intraspecific variation may also be important in allowing species to track environmental variation and in promoting trait convergence to an environmental optimum (Henn et al., 2018). However, other studies have suggested that intraspecific trait variation responds differently to environmental gradients compared to interspecific variation (Laughlin et al., 2017). It therefore remains unclear how intraspecific trait shifts and abundance shifts affect community weighted mean traits following changes in environmental conditions (Roscher et al., 2018), such as increased nitrogen levels.

Nitrogen enrichment can directly and indirectly affect functional composition. Increases in soil nitrogen levels will typically directly increase community SLA (and decrease LDMC) by favouring the establishment and growth of faster growing species (Wright et al., 2004), but also because individual plants produce leaves with higher SLA (Shipley & Almeida-Cortez, 2003; Siefert & Ritchie, 2016). Nitrogen enrichment could also indirectly affect functional composition, in particular, by causing a loss of plant diversity and changes in consumer abundance. A loss of species richness might reduce intraspecific SLA values due to lower light competition in open, species poor communities (Lipowsky et al., 2015). A change in the abundance of foliar fungal pathogens with nitrogen enrichment (Dordas, 2008) might reduce SLA if pathogens reduce the relative abundance of fast growing species or cause species to shift trait expression towards lower SLA, due to the lower investment in defence against pathogens by fast species compared to slow species (Blumenthal, Mitchell, Pysek, & Jarosík, 2009; Cappelli, Pichon, Kempel, & Allan, 2020). The responsiveness of functional composition to environmental changes could also differ between fast and slow communities, as high SLA species, which are more competitive for light, might show higher plasticity and intraspecific variation in SLA (Crick & Grime, 1987; Freschet et al., 2015). However, the amount of variation in SLA along the leaf economics spectrum is not well known. It is not known whether various environmental factors alter functional composition principally through favouring species with certain traits (abundance shifts) or by causing intraspecific shifts in traits.

Shifts in community functional composition can have large effects on a range of ecosystem functions (Díaz et al., 2007; Lavorel & Grigulis, 2012; Allan et al., 2015; Ratcliffe et al., 2017; Cappelli et al., 2020; Pichon, Cappelli, Soliveres, Mannall et al., 2020; Pichon, Cappelli, Soliveres, Hölzel et al., 2020). However, many studies looking at the effects of functional trait composition on ecosystem function calculate trait shifts using literature values, or measure a single mean value per species, i.e. ignoring intraspecific variation (see Violle et al., 2012). It is now known that intraspecific shifts can explain a large proportion of the variation in community mean traits, however, the importance of intraspecific changes in affecting ecosystem functioning is still poorly known and under debate (Albert et al., 2011). Intraspecific trait changes might not have the same effect on ecosystem functioning as interspecific shifts because the strong correlations between leaf economic traits break down within communities (Messier, McGill, Enquist, & Lechowicz, 2017; Anderegg et al., 2018), which could be due to lower variation in leaf life span within communities (Funk & Cornwell, 2013) or different correlations between traits, within and across species (Laughlin et al., 2017). It is also possible that response and effect trait correlations change within species compared to across species, i.e. that intraspecific trait responses to environmental factors are decoupled from their effects on function, however, this has not been tested. It is crucial to understand the extent to which intraspecific variation simultaneously contributes to explaining trait responses to the environment and trait effects on functioning, if we are to predict how environmental change alters ecosystem functioning.

To quantify the extent of intraspecific shifts in response to the direct and indirect effects of N, and the consequences of these shifts for ecosystem functioning, we conducted a field experiment manipulating N enrichment, species richness, functional composition and fungicide application. We therefore manipulated direct effects of N and two indirect effects (loss of diversity and changes in consumer communities). We further, independently manipulated the mean SLA of our communities by establishing plots with different combinations of high and low SLA species, which also created a large gradient in initial LDMC. Communities were initially sown with species at equal abundance, but abundances and intraspecific trait expression could shift freely within and between communities, allowing us to assess trait convergence and divergence from a known starting point. We could therefore quantify the contribution of shifts in abundance and intraspecific trait expression to changing community mean traits, how large this variation was relative to the compositional variation between plots, and whether the degree of intraspecific and abundance shifts depended on the initial (sown) trait composition. We fitted two structural equation models for SLA and LDMC, first to quantify the intraspecific and abundance shifts, from the initial trait composition, due to N enrichment, fungicide and species richness, and how these shifts contributed to total variation in community weighted mean traits. We then fitted a second pair of SEMs linking intraspecific and abundance shifts to above and belowground biomass production, instead of the community weighted mean traits, to test the effect of intraspecific trait shifts on biomass production. Previous studies analysing effects of trait shifts on function calculated changes in biomass for individual species (Liancourt et al., 2015), or compared models fitting community weighted mean calculated using traits of different origins (Roscher et al., 2018). Our approach has the advantage of simultaneously estimating the relative response and effect of each component of the traits shifts.

Within this framework, we tested the following hypothesis:

1. Intraspecific shifts in trait expression and shifts in the relative abundance of species are similarly important in determining functional composition, i.e. the overall variation in community weighted mean traits between communities.
2. N enrichment, species richness and fungicide application all favour fast-growing species, i.e. they increase mean SLA and decrease mean LDMC.
3. Both intraspecific shifts in trait expression and shifts in relative abundances affect ecosystem functioning, and the relative importance of intraspecific shifts in driving responses and effects is similar

## Material and Methods

### Field site

We conducted the study in the PaNDiv experiment, a large grassland experiment located in Münchenbuchsee, Switzerland. The PaNDiv experiment started in autumn 2015 and manipulates plant functional composition, species richness, N enrichment and fungicide application in a full-factorial design. We assembled plant communities using a pool of 20 species common in central European grasslands. We divided the species into two groups according to their SLA and leaf N content. The two groups contained both herbs and grasses and were classified as having a slow growing (low SLA, low N content), or fast growing strategy (high SLA, high N). Experimental communities contained one, four or eight species and at the four and eight species levels, communities contained either slow or fast growing species, or a mix of both (monocultures could contain only one functional strategy). The species in each community were randomly drawn from the particular species pool (i.e. all species, slow or fast growing), for a total of 50 different combinations. This created a large gradient in mean SLA values (15 to 29 m^2^ kg^-1^) and mean LDMC values (208 to 290 mg g^-1^) between communities, which is comparable to trait values along a central European land use intensity gradient (SLA: 13-32 m^2^ kg^-1^, LDMC: 220-420 mg g^-1^, Breitschwerdt, Jandt, & Bruelheide, 2018). Each combination of species was grown four times; in control conditions, with N enrichment (100 kg N ha^-1^ y^-1^ as urea twice a year in April and June), with fungicide application (“Score Profi”, 24.8 % Difenoconazol 250 g.L^-1^, four times during the growing season) and with both N and fungicide together, resulting in 200 plots. Communities were sown with equal numbers of seeds per species, corrected for germination rates (total 1000 seedlings m^-2^), on 2×2m plots separated by a 1m path. The plots were randomly distributed in four blocks, each containing every species combination, each time with a randomly assigned treatment.

The site was weeded and ploughed before the start of the experiment. To maintain the different diversity levels, we weeded each plot three times per year, in April, July and September, enabling us to keep total weed abundance to only 5% of total cover, on average. The whole field was mown in mid-June and late August and the biomass was removed, which simulates extensive to intermediate grassland management in the Swiss lowlands. The carbon and nitrogen organic content were measured in June 2019: 2.35-4.70 % organic C and 0.25-0.43% organic N (Walde et al., In prep.). The mean volumetric water content of 5.95% indicated rather dry conditions when we collected the samples for the present study (mean from June to August 2017). Further information about the design of PaNDiv and field characteristics can be found in Pichon, Cappelli, Soliveres, Hölzel et al. (2020).

We harvested above ground biomass production before mowing in August 2017 on two quadrats of 20×50 cm in the centre of each plot, clipping vegetation above 5 cm. The samples were dried at 65°C for 48h and weighed. Percentage cover of target species, weeds and bare ground, was recorded at the same time. Aboveground biomass production was corrected for weed cover by multiplying the biomass by the proportion of target (non-weed) species. Belowground biomass was measured in autumn 2017 by taking two cores per plot to 20 cm depth (440 cm^3^ of soil). We homogenised the two samples and used a subset of 40g fresh soil in which we washed and sorted out the roots. We then dried the roots at 65°C for 48 hours. To calculate belowground biomass per g of dry soil, we estimated soil bulk density by weighing 40g of soil from the same plot before and after drying for 24 hours at 105°C. Aboveground biomass data were square-root transformed and belowground biomass data were log-transformed in order to meet model assumptions (see analysis section).

### Leaf sampling

We collected leaf samples on the 200 plots over two weeks in August 2017, taking five leaves per species per plot. Specific leaf area and leaf dry matter content were measured following the protocol of Garnier, Shipley, Roumet, and Laurent (2001). We measured leaf fresh weight and recorded the leaf area with a leaf area meter (LI-3000C, LI-COR 253 Biosciences), after overnight rehydration in deionised water in the dark. We dried the samples at 65°C for two days and measured their dry weight. We calculated SLA (area/dry weight) and LDMC (dry/fresh weight) per species per plot by averaging the values of the five samples. Two species, *Anthriscus sylvestris* and *Heracleum sphondylium* established very slowly in the experiment and were rare in the first years. Because of a lack of plants of these two species, we excluded them completely from the community trait calculations.

In order to characterise community functional composition and compare the changes in traits due to shifts in species relative abundance or in intraspecific trait values, we calculated four community trait metrics (Figure 1). 1) The **sown trait** value per 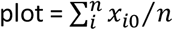; with *x*_i0_ being the trait value of species *i* in the control monocultures and n the number of species per plot. We used this as the baseline measure, as it indicates the trait value in control (unfertilised, no fungicide) conditions, for species experiencing intraspecific competition only. 2) The Δ **abundance shift** per 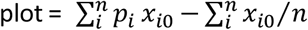; with *p*_i_ being the relative abundance of species *i* in a given plot, and *x_i0_* the trait value of species *i* in the control monoculture. 3) The Δ **intraspecific shift** per 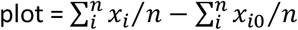; with *x*_i_ being the trait value of the species *i* measured in a particular plot and n the number of species in a given plot. 4) The overall **community weighted mean** 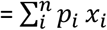; with *p_i_* the relative abundance of the species *i* and *x_i_* the trait value of species *i* per plot. We did not include monocultures in the analysis, as the Δ values were equal to 0, by definition, in a monoculture.

**Figure 1:**
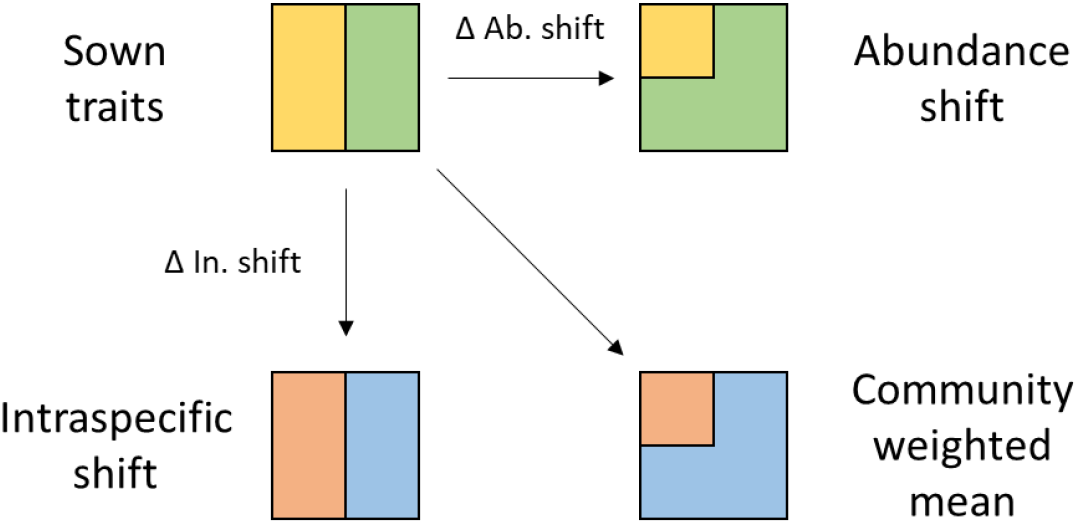
Schematic description of the components of community weighted mean traits, for a community of two species, represented by coloured rectangles. Shifts in community weighted mean traits are due to shifts in species relative abundances (Δ ab. shift, represented by a change in the size of the rectangles) and to intraspecific trait shifts (Δ in. shift, represented by a change in colours).

Seven species did not establish or were present at very low percentage covers in 36 plots, and traits of these species could not be sampled (44 of 716 data points). Although this did not influence the abundance shift and community weighted mean calculations, we replaced these values with monoculture data in the intraspecific shift calculations and therefore made the conservative assumption that the intraspecific shift would have been 0 for these species. However, there was no difference in the intraspecific shift effect, if these species were completely excluded, see Table S1.

### Analysis

We tested how our different factors influenced the variation in community mean SLA and LDMC, and how the variation in these traits affected ecosystem functioning, in a two-step procedure. First, we fitted a set of linear mixed effect models to screen for potential interactions, and second, a set of structural equation models (Grace, 2006) to partition the contributions of abundance and intraspecific shifts to community weighted mean traits and ecosystem functioning. All analysis were conducted in R (Bates, Mächler, Bolker, & Walker, 2015; R Core Team, 2019). Interactions between all of the experimental treatments are plausible, however, incorporating large numbers of interactions in SEMs is challenging. As there is no theory to predict which interactions should be most important, we used linear mixed effects models to determine which interactions to include in the SEMs. We fitted four linear mixed models for the two abundance shifts (Δ abundance shift in SLA and LDMC) and two intraspecific shifts (Δ intraspecific shift in SLA and LDMC). Each model included the effect of nitrogen enrichment, fungicide, plant species richness, sown mean traits, and all two-fold interactions, as fixed effects and block and species composition (the randomly assembled sets of species) as random terms. The data were not transformed because the errors were normally distributed and the variance homogenous. We simplified the initial models using likelihood-ratios to drop terms that did not significantly improve overall model fit (Zuur, Ieno, Walker, Saveliev, & Smith, 2009).

We then fitted two different structural equation models for each trait. We first tested the relative importance of intraspecific and abundance shifts in determining community weighted mean traits, and second tested the effect of the two shifts on ecosystem functions. In the first set of models, we fitted direct paths from nitrogen enrichment, species richness, fungicide and sown traits to the Δ abundance shift and the Δ intraspecific shift, with the hypothesis that nitrogen, richness and fungicide would increase the two Δ SLA shifts and decrease the two Δ LDMC shifts. We added paths from interactions between sown SLA and the three other factors to intraspecific shift SLA, to test how the treatment effects depended on the community growth strategy, and from an interaction between richness and fungicide to abundance shift SLA, as these interactions were significant in the LMMs (Table S2). For the LDMC model, we added an interaction between richness and nitrogen on abundance shift LDMC. To assess the relative importance of sown community composition, abundance shifts and intraspecific shifts in determining overall functional composition, we then fitted direct paths from the sown trait values, and the two Δs, to the community weighted mean trait. The second set of SEMs tested the relative importance of the treatments and aspects of functional composition in affecting above-and belowground biomass. The structure was similar, but we fitted paths to above-and belowground biomass rather than community weighted mean traits. We added paths from nitrogen, fungicide and species richness to aboveground and belowground biomass, to include additional direct (hypothesised positive) effects of these factors on biomass, not occurring through changes in SLA or in LDMC. We deliberately chose to fit CWM traits and biomass values in two separate models to more clearly show the factors affecting each of them. We do not want to test for the overall effect of CWM traits on biomass because we aim to test the relative importance of intraspecific and abundance shifts in separately determining functional composition (CWM) and in affecting biomass production. However, fitting a model with both CWMs and biomass together does not affect the results.

## Results

### Trait responses to the experimental treatments

We observed large intraspecific trait and abundance changes, which altered community weighted mean trait values. Community weighted mean (CWM) values of SLA and LDMC were affected by compositional variation (the sown community values) but also by intraspecific and abundance shifts (Figure 2 and Table S3 and S4). For both traits, intraspecific shifts contributed as much to variation in the final CWM traits as abundance shifts, which indicates substantial intraspecific variation in these two plant traits. For LDMC the intraspecific shifts even had a similar magnitude of effect on the CWM as sown compositional variation between the plots (path coefficient of 0.6±0.05 for sown LDMC and 0.53±0.05 for intraspecific shifts), while for SLA the sown compositional variation had the same effect as intra and abundance shifts together (path coefficient for sown SLA of 0.98±0.04, intraspecific shift 0.44±0.03 and abundance shift 0.54±0.04). Compositional variation, intraspecific and abundance shifts combined to explain the majority of the variation in CWM traits (88% for SLA and 75% for LDMC), however, the R^2^ values for the CWMs were <1 in the SEMs. This is partly because intraspecific and abundance shifts did not have an entirely additive effect on the CWMs. However, including an interaction between Δ interspecific and Δ abundance shift only slightly increased the R^2^ (from 0.876 to 0.885 for SLA, and from 0.754 to 0.755 for LDMC, data not shown). The remaining variation could be due to non-linear effects of the different factors or other, possibly higher order interactions between composition variation and the intraspecific and abundance shifts.

**Figure 2:**
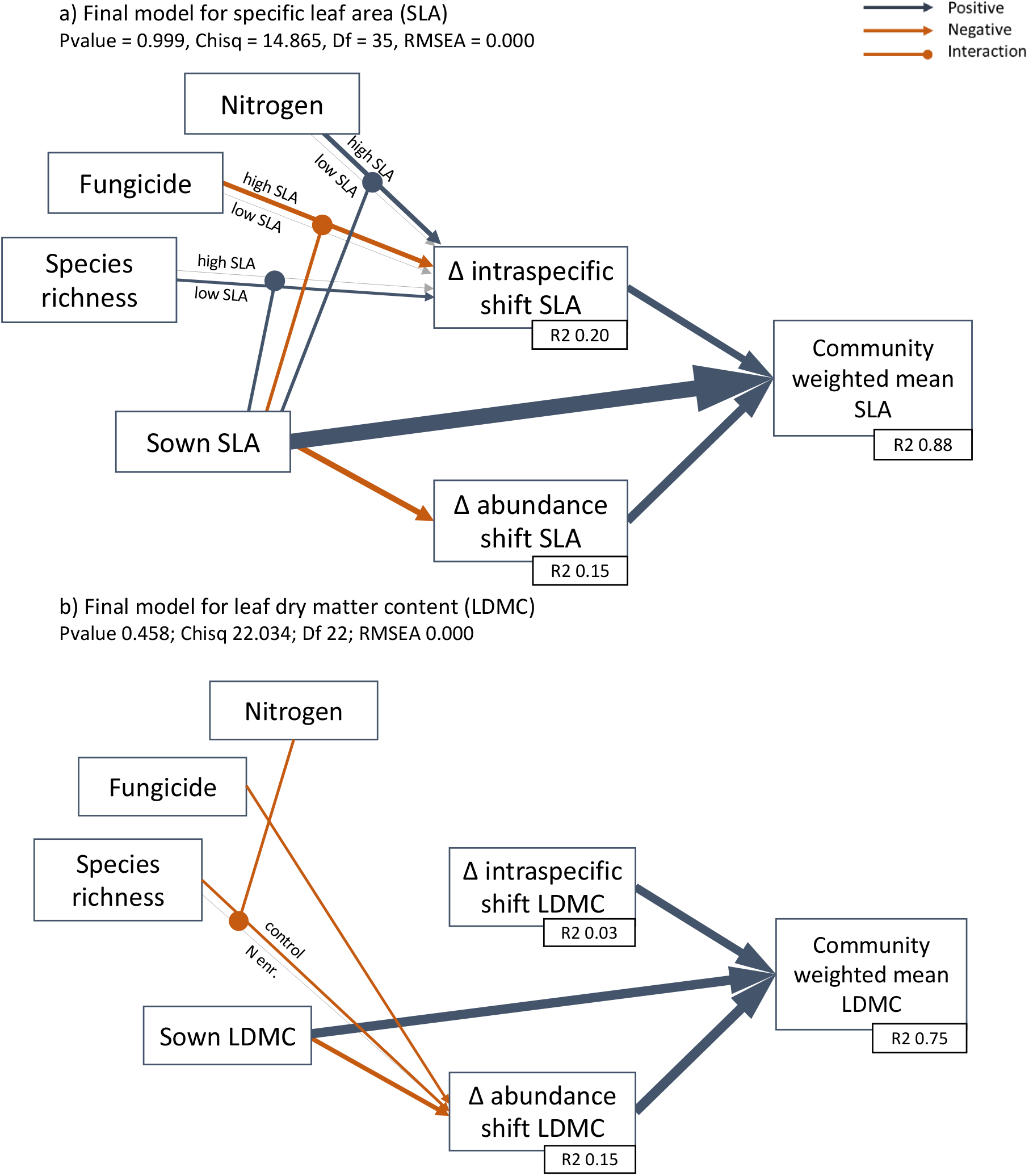
Effect of nitrogen, fungicide, species richness and sown mean traits on community weighted mean trait values through a shift in species abundance (Δ abundance shift) or in intraspecific trait values (Δ intraspecific shift). Significant output of structural equation models for a) specific leaf area (Table S3) and b) leaf dry matter content (Table S4). N = 120 plots. Blue arrows show a positive effect, red a negative effect. The size of the arrows is proportional to the path coefficient. The effect of nitrogen, fungicide and species richness on the intraspecific shift in SLA depended on the initial sown community, and the effect of richness on abundance shift LDMC depended on nitrogen. These interactions are represented here by round-headed arrows and two lines, one showing the effect of, e.g., N addition at high vs. low SLA. Interactions are also plotted as partial plots in Figure 3.

Plant species richness, nitrogen addition and fungicide application, in interaction with sown SLA, all caused intraspecific shifts in SLA (Figure 2a) but had no effect on intraspecific shifts in LDMC (Figure 2b). In mixed plots with low sown SLA, species, on average, increased their SLA compared to monoculture, and the nitrogen and fungicide treatments did not cause intraspecific shifts in SLA in these plots (Figure 3a and c). In plots with high sown SLA, species increased their SLA only when N was added, and otherwise they decreased their SLA (Fig. 3c). Fungicide had the opposite effect and SLA was reduced when fungicide was added to plots sown with high SLA species (Fig. 3a). Sown diversity also changed the response: in plots with four species, SLA responses were very variable whereas in plots sown with 8 species, SLA increased in plots sown with low SLA species, and declined in plots sown with high SLA species (Figure 3b).

**Figure 3:**
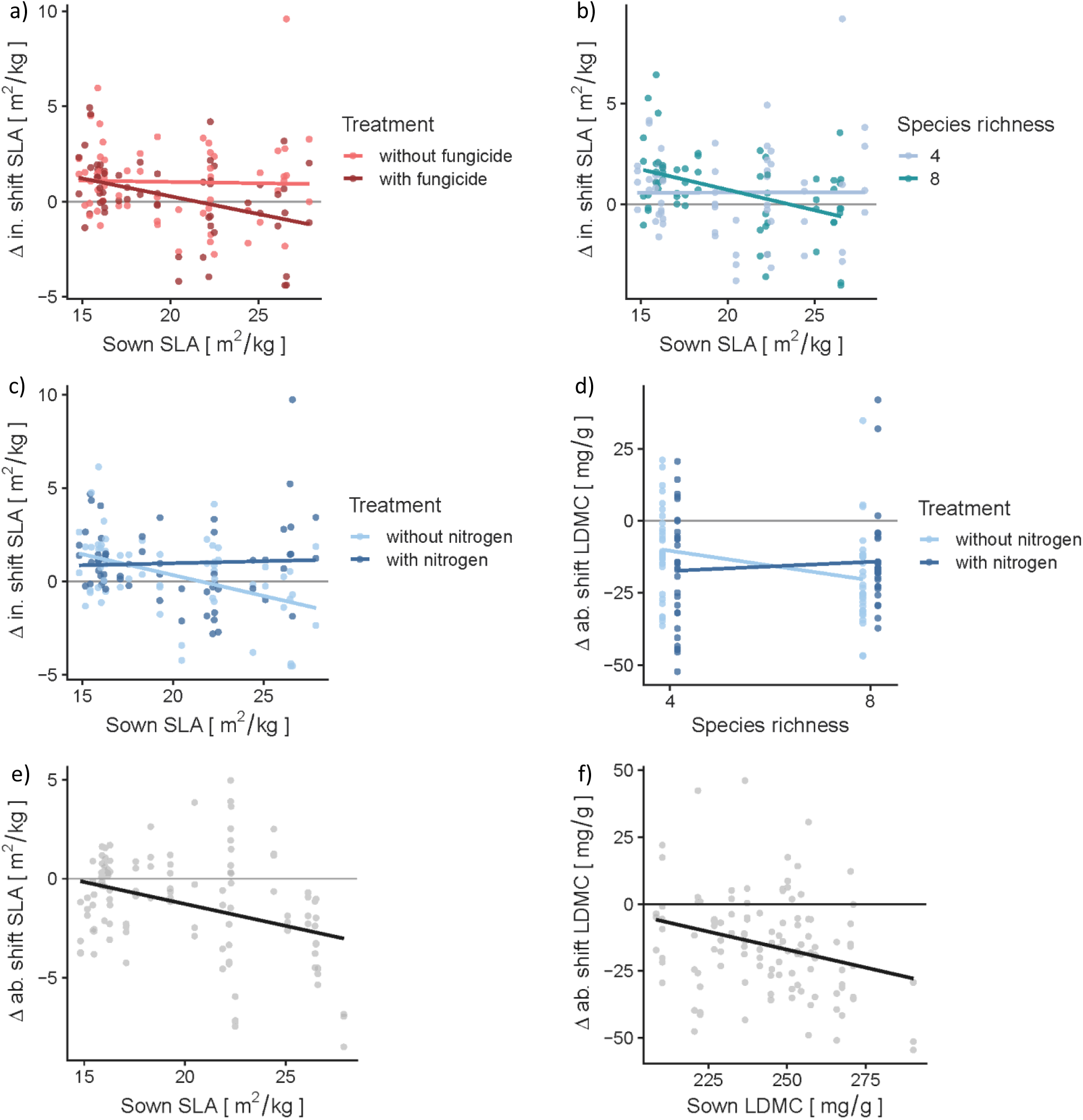
Partial plots visualising the structural equation models output shown in Figure 2. Effect of sown specific leaf area (SLA) on the difference between intraspecific shift and sown SLA (Δ in. shift SLA) depending on a) fungicide, b) species richness and c) nitrogen. d) Effect of species richness on the difference between abundance shift and sown LDMC (Δ ab. shift LDMC) depending on nitrogen enrichment. Effect of sown trait on the difference between abundance shift and sown trait (Δ ab. shift) e) for SLA and f) for LDMC. The 0 line indicates no changes from the sown values. X-axis units are back-transformed values, y-axis are back-transformed residuals of the target explanatory variables on the remaining explanatory variables. N = 120 plots.

Abundance shifts tended to decrease mean SLA and LDMC of communities, particularly at high sown values of the traits (Figure 3e-f), indicating that low SLA and LDMC species dominated the experimental communities. Communities sown with generally high SLA species became more strongly dominated by species with a lower SLA, leading to a reduction in CWM SLA in communities with high sown SLA (Figure 4a-b). In communities sown with high LDMC species, it was also the low LDMC species that dominated (Figure 3f, and 4c-d). Whereas the abundance shifts that altered SLA could not be explained by any of the other treatments, fungicide application led to abundance shifts that reduced mean LDMC (Figure 2). This indicates that species with a relatively low LDMC dominated in plots where foliar fungal pathogens were removed. Species richness and nitrogen also interacted to affect abundance shifts in LDMC: low LDMC species dominated more in 8 species plots but N dampened this effect (Figure 3d).

**Figure 4:**
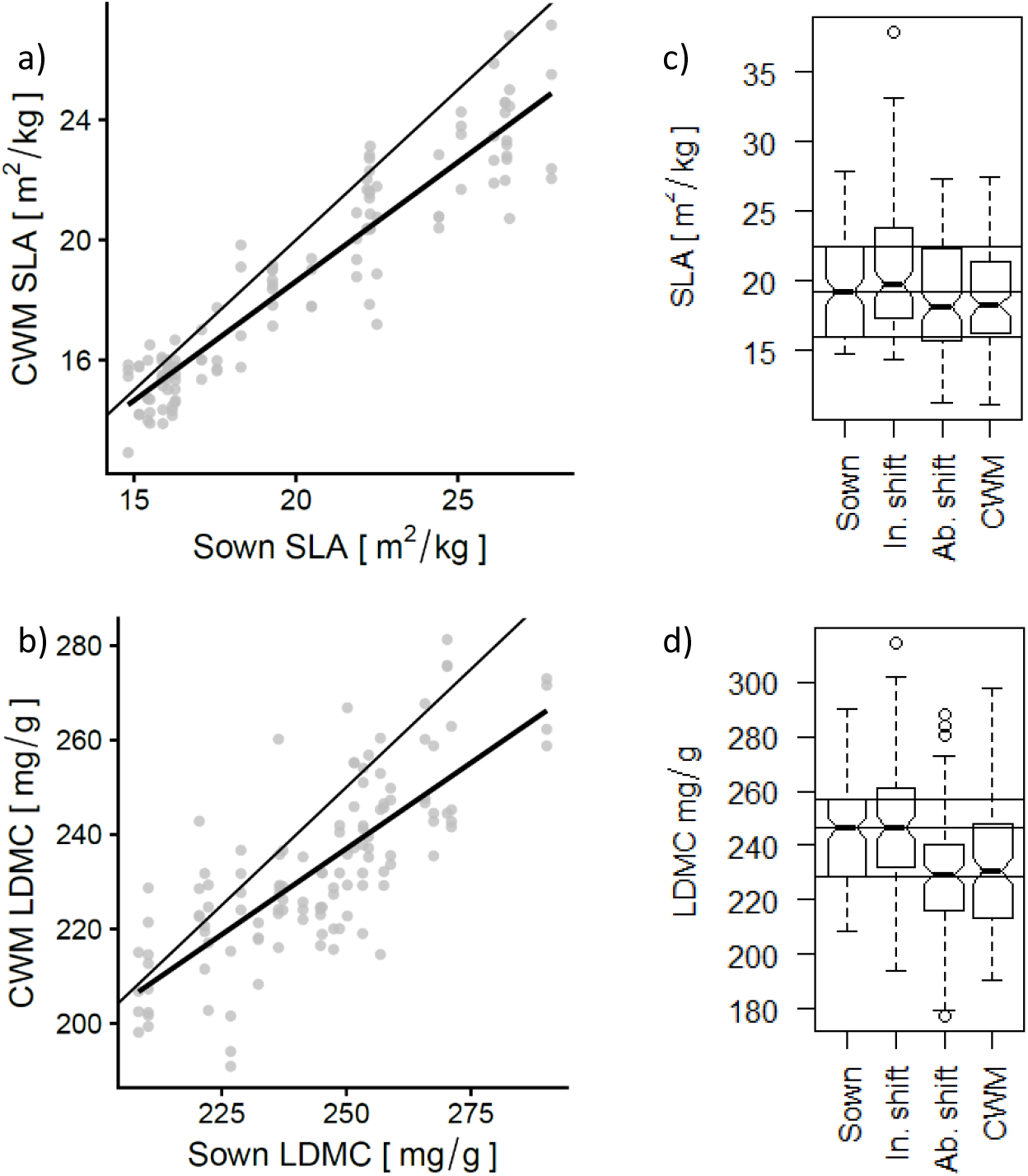
Effect of sown traits on community weighted mean (CWM) trait values, **a)** for specific leaf area (SLA) and **b)** for leaf dry matter content (LDMC). The 1:1 line is pictured in light grey. Partial plots visualising the structural equation models output are shown in Figure 2. X-axis units are back-transformed values, y-axis are back-transformed residuals of the target explanatory variables on the remaining explanatory variables. The amount of variation in community mean (**c**) SLA and (**d**) LDMC is represented by the adjacent boxplots, showing the difference in community mean when calculated using sown values, intraspecific shift, abundance shift or community weighted mean values. N = 120 plots.

Overall, intraspecific and abundance shifts had opposite effects on community mean trait values, by leading to increases and decreases in SLA and LDMC. In addition, the intraspecific and abundance shifts resulted in a convergence of the communities towards intermediate values of CWM SLA and LDMC by slightly increasing mean trait values in plots sown with low SLA/LDMC species and reducing them in plots sown with high SLA/LDMC species (Figure 4a-b).

### Effects of trait shifts on ecosystem function

Whereas intraspecific trait shifts explained the same amount of variation in CWM traits as abundance shifts, this did not translate into an effect of intraspecific shifts on ecosystem functioning (Figure 5-6 and Table S5-6). Only abundance shifts affected above and belowground biomass. Increased abundance of high SLA species increased aboveground biomass production and decreased belowground production, indicating contrasting effects of SLA on these two ecosystem functions (Figure 6a-b). Compositional variation was also important as a high sown SLA further decreased belowground biomass, although sown SLA had no effect on aboveground biomass production, after taking the abundance shifts into account. Similarly, LDMC shifts only affected functioning through shifts in abundance. Shifts in species abundances towards higher mean LDMC increased belowground biomass (Figure 6c-d), which agrees with the SLA effect and indicates that plots dominated by slow growing species had higher root production. High values of sown LDMC also increased belowground biomass. None of the measures of LDMC affected aboveground biomass and in the LDMC SEM, aboveground biomass was only directly affected by nitrogen enrichment. Species richness had no direct effect on functioning, however, this is probably because we only included data from the 4 and 8 species plots, which means there was very low variation in species richness.

**Figure 5:**
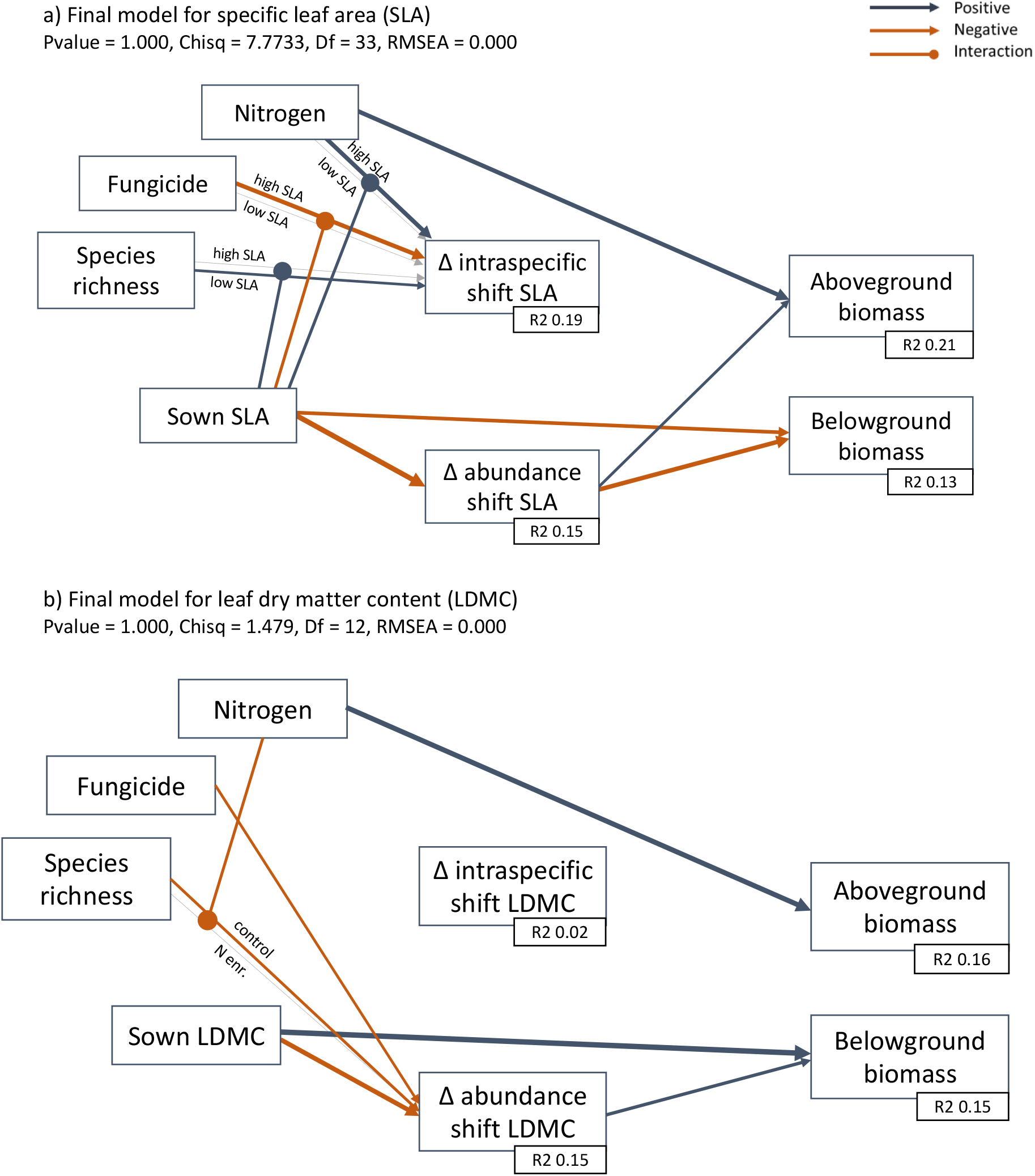
Effect of nitrogen, fungicide, species richness and mean sown traits on aboveground and belowground biomass. Effects are direct or through a shift in species abundances (Δ abundance shift) or intraspecific trait values (Δ intraspecific shift). Significant output of structural equation models for a) specific leaf area (Table S5), and b) leaf dry matter content (Table S6). N = 120 plots. Blue arrows show a positive effect, red a negative effect. The size of the arrows is proportional to the path coefficient. The effect of nitrogen, fungicide and species richness on the intraspecific shift in SLA depended on the initial sown community, and the effect of richness on abundance shift LDMC depended on nitrogen. These interactions are represented here by round-headed arrows and two lines, one showing the effect of, e.g., N addition at high vs. low SLA. Interactions are also and are plotted as partial plots in Figure 3.

**Figure 6:**
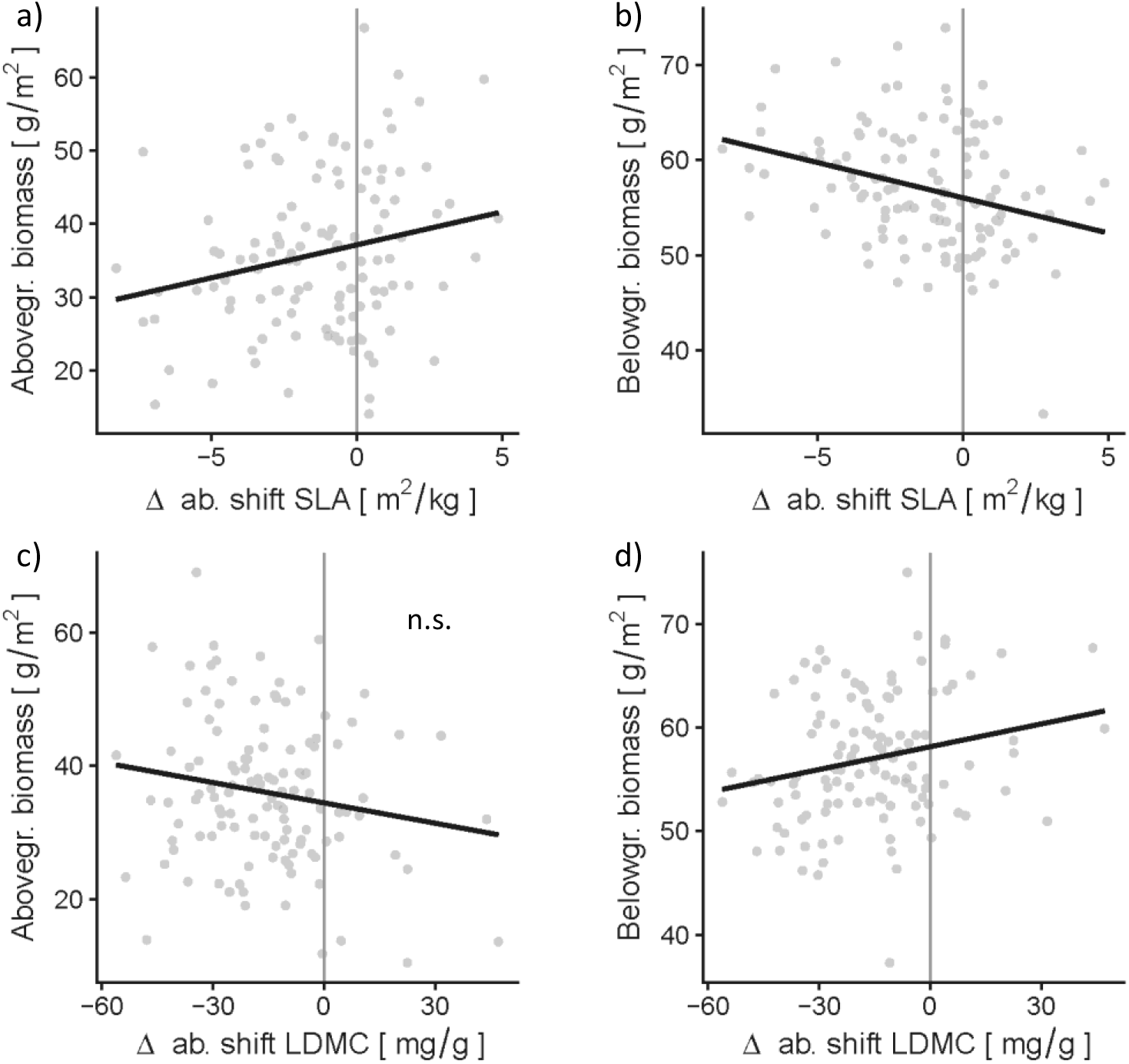
Partial plots visualising the structural equation models output of Figure 5. Effect of the difference between abundance shifts and sown specific leaf area (Δ ab. shift SLA) on **a)** aboveground biomass and **b)** belowground biomass. Effect of the difference between abundance shifts and sown leaf dry matter content (Δ ab. shift LDMC) on **c)** aboveground biomass (non-significant) and **d)** belowground biomass. The 0 line indicates no changes from the sown values. X-axis units are back-transformed values, y-axis are back-transformed residuals of the target explanatory variables on the remaining explanatory variables. N = 120 plots.

## Discussion

### Intraspecific trait shifts have large effects on overall functional composition

Our results showed that changes in functional composition due to abundance and intraspecific shifts were of similar importance in explaining overall community weighted mean trait measures. This agrees with an increasing volume of literature on the importance of intraspecific variation in SLA (Albert et al., 2011; Lepš et al., 2011; Violle et al., 2012; Carmona, de Bello, Mason, & Lepš, 2016), and highlights substantial variation also in LDMC, which is often considered less plastic than SLA (Garnier, Shipley et al., 2001). The large contribution of intraspecific shifts, relative to abundance shifts, to overall community weighted means is perhaps not surprising, given that all plots were located within the same field (Cordlandwehr et al., 2013; Petruzzellis et al., 2017). However, we experimentally created a large gradient in mean leaf traits, similar to the variation between extensively and intensively managed grasslands (Breitschwerdt et al., 2018), by deliberately selecting a pool of species to cover a large range in SLA. Even relative to this large initial variation in trait composition, intraspecific trait shifts explained a substantial amount of the total functional variation between communities. In addition, applying nitrogen and fungicide treatments meant that there was substantial variation in resources and enemy pressure between plots. This large contribution of intraspecific variation to overall community mean traits therefore challenges the idea that within species trait shifts only affect functional composition when environmental conditions and species are similar.

The abundance and intraspecific shifts caused large, and opposing, changes in functional composition relative to the originally sown communities. Shifts in species abundances decreased both mean SLA and LDMC across the whole field. In our experiment, grasses and herbs differed strongly in LDMC so that high mean LDMC indicates a larger relative cover of grasses (Figure S1). The decrease in both SLA and LDMC due to abundance shifts therefore indicates that slow growing herbs (low SLA, low LDMC) dominated the experimental communities. In contrast, intraspecific trait shifts (from monocultures to mixtures) increased SLA in both slow and fast dominated communities. Freschet et al. (2015) hypothesised that acquisitive (fast) species should be more plastic than conservative (slow) ones, however, we find that slow dominated communities could also shift their SLA values substantially. Intraspecific shifts towards higher SLA in mixed communities could be driven by higher light competition compared to monocultures (Lipowsky et al., 2015). However, the intraspecific shifts in SLA were not constant across plots and were modulated by N enrichment, fungal pathogen presence and species richness, see below. Opposing intraspecific and abundance shifts and opposing trait shifts in fast and slow species, led to an overall convergence in community weighted mean SLA towards intermediate values, perhaps reflecting the fairly fertile but water limited conditions on our field site. These results show that intraspecific trait shifts can cause large changes in functional composition but that changes due to shifts in abundance and intraspecific trait values can be decoupled. This could be due to correlations between functional traits breaking down at the intraspecific scale (Laughlin et al., 2017). Low SLA species might be more resistant to low water availability compared to high SLA species (Poorter, Niinemets, Poorter, Wright, & Villar, 2009), partly because they have particular root traits (leading to higher root production, Figure 5a) and might therefore be favoured on our site. However, at the same time light competition in mixed communities favours high SLA values (Lipowsky et al., 2015). If SLA and root traits are decoupled at the intraspecific level, then individuals might increase their SLA in response to shading (leading to positive intraspecific shifts), while at the same time low SLA species are favoured because they are better at competing for soil resources, leading to opposing intra and interspecific trait shifts. Such opposing responses were also found by Kichenin, Wardle, Peltzer, Morse, and Freschet (2013) along an elevational gradient. They showed that a correlation between SLA and leaf pubescence at the interspecific level broke down within species, so that at high elevation the species tended to produce lower SLA leaves to cope with water scarcity but at the same time high SLA species increased in abundance because their pubescent leaves had better water retention capacity. Contrasting intraspecific and abundance shifts show that within and between species trait variation may respond to different aspects of the environment, suggesting that response traits could differ within and between species.

### The environmental conditions causing trait shifts

Manipulating environmental conditions changed community SLA through intraspecific shifts, and LDMC through abundance shifts. These opposing patterns could be due to lower correlations between leaf traits under the different treatments. Laliberté and Tylianakis (2012), showed that an increase in community SLA following nutrient enrichment was linked to a decrease in leaf thickness, therefore leading to higher SLA without changing LDMC values. The different ways in which environmental conditions affect community SLA and LDMC could therefore lead to a local decoupling of these two functional traits (Laughlin et al., 2017; Messier et al., 2017; Anderegg et al., 2018). These results further emphasise that intraspecific variation in traits may drive different environmental responses compared to interspecific variation and that trait correlations can break down at certain scales.

Exclusion of fungal pathogens affected intraspecific shifts in SLA. In communities sprayed with fungicide, SLA values towards intermediate values, due to an increase in slow communities and a decrease in fast communities. The intraspecific shifts reflected the change in SLA from monocultures to mixtures and therefore the trait changes associated with a shift from intra to interspecific competition. Interspecific competition could lead to either trait divergence when traits are linked to niche differences, or trait convergence when they are linked to competitive ability differences (Mayfield & Levine, 2010) and previous studies have shown that SLA is linked to competitive ability differences, rather than niche, differences (Kraft, Godoy, & Levine, 2015). The presence of fungal pathogens can reduce competitive interactions between plants (Chesson, 2000; Mordecai, 2011) and removing pathogens might therefore have increased interspecific competition in our communities and led to convergence in SLA.

Fungicide spraying increased the abundance of low LDMC species. This result is in line with the growth-defence trade-off (Heckman, Halliday, & Mitchell, 2019), which would predict that fungicide favoured the establishment of fast growing species (low LDMC) because their energy was invested more in growth than in defence. We may have observed this abundance shift in LDMC not in SLA because LDMC reflects structural components of the leaves that contribute to physical defence against herbivores and pathogens (Ibanez, Lavorel, Puijalon, & Moretti, 2013; Descombes et al., 2017). Fungicide therefore decreased both SLA and LDMC but due to intraspecific shifts and abundance shifts respectively.

Nitrogen addition led to intraspecific shifts in SLA but did not favour high SLA species in general. N enrichment directly increased above ground biomass, showing that productivity is N limited at our site, however it did not increase the abundance of high SLA species, contrary to our expectations (Laliberté, Shipley, Norton, & Scott, 2012; Lavorel & Grigulis, 2012). N enrichment may not have favoured fast, high SLA species, because these remained limited by water or other nutrients, and therefore SLA was not the key trait determining response to N alone (Firn et al., 2019). Dry conditions in summer could have prevented fast growing species from taking advantage of the N addition and increasing their abundance (Poorter et al., 2009; Jung et al., 2014; Rosbakh, Römermann, & Poschlod, 2015). However, nitrogen enrichment did cause intraspecific trait shifts and affected the degree of trait convergence between communities. Nitrogen only increased SLA in fast growing communities, in slow growing communities SLA increased regardless of whether communities were fertilised or not. Nitrogen addition therefore reduced the convergence in SLA between communities and led to general increases in SLA, as expected. This suggests that fast species reduced their SLA when growing in mixtures compared to monocultures, possibly due to increased water or nutrient competition (Poorter et al., 2009), but that N enrichment dampened this effect.

We found that trait convergence occurred only in eight species communities, not in plots with four species. Lipowsky et al. (2015) also found greater intraspecific shifts in SLA at higher species richness. The greater convergence at higher species richness could have been caused by greater trait variation and therefore more opportunities for trait shifts in eight species communities. This would be a type of sampling effect, where eight species communities are more likely to contain species with high genetic variation in traits or with high trait plasticity. The greater convergence in SLA in species rich communities supports the idea that diverse communities are better able to shift their traits to cope with environmental variation (Vogel et al., 2019).

Intraspecific shifts in LDMC were also large: they contributed as much to variation in community mean traits as compositional differences, but we could not explain them. Intraspecific shifts in LDMC could have been driven by variation in microclimatic conditions, e.g. water availability, between plots (Jung et al., 2014) or by random processes leading to differential establishment of genotypes in the plots. However, it seems unlikely that genetic variation alone could explain such large intraspecific variation, and different environmental factors driving different plastic shifts in the various plots might be a more plausible explanation (Siefert et al., 2015; Lajoie & Vellend, 2018).

### Effect on biomass

Although intraspecific shifts in SLA and LDMC had large effects on community weighted mean values, this did not translate into a change in above or belowground biomass production. This result runs counter to the study by Liancourt et al. (2015) which showed that plastic shifts in SLA affected biomass production. However, that study looked at changes in biomass production for individual species under different treatments, which makes it difficult to compare with our measure of overall community biomass production. However, our results support those of Roscher et al. (2018) who showed no significant difference in model fits when calculating the effect of CWM traits on biomass using traits measured in monocultures or in mixtures. Intraspecific trait shifts may not have affected functioning because variation in leaf traits is not linked to variation in resource use within species (Messier et al., 2017; Anderegg et al., 2018). The leaf economics spectrum results from a trade-off between high photosynthetic rates in leaves with a large area per unit of invested mass, (high SLA; the acquisitive strategy) vs. a long leaf lifespan and thicker leaves (low SLA; the conservative strategy) (Wright et al., 2004). Funk and Cornwell (2013) showed that the link between leaf traits and resource use strategies depends on the amount of variation in leaf lifespan within a community, meaning the relationship would break down at a small scales. However, for SLA the intraspecific shifts in response to the experimental treatments, show that intraspecific responses are broadly in line with expectations, although they are more complex than hypothesised. However, these responses seem to be decoupled from the effects of intraspecific trait changes on ecosystem functioning. This may indicate that response-effect trait correlations are different within vs. between species. We show here that intraspecific changes in SLA did not affect functioning, suggesting that even when large, intraspecific trait variation is not necessarily important for ecosystem functioning.

Root biomass production increased in communities dominated by slow species. Communities sown with low SLA and low LDMC species, and those in which low SLA and low LDMC species increased their abundance, produced most root biomass. The effect of SLA on root biomass production is rarely assessed (Mommer & Weemstra, 2012; Bergmann, Ryo, Prati, Hempel, & Rillig, 2017) but we observe that slow growing species invested more in roots than fast growing ones. Plants adapted to resource poor environments would be expected to invest more resources belowground, as this would give them an advantage under dry or low nutrient conditions (Freschet et al., 2015). In addition, the joint effects of SLA and LDMC on root production indicate that it is slow growing grasses (low SLA and high LDMC) that had highest root biomass (Gastine, Scherer-Lorenzen, & Leadley, 2003; Ravenek et al., 2014). Therefore, an overall reduction in community SLA and LDMC due to abundance shifts had contrasting effects on belowground biomass production.

Aboveground biomass production was altered by changes in SLA but shifts in abundance and compositional variation had opposing effects. Where plots became dominated by fast (higher SLA) species, above ground biomass production increased. However, sown community SLA had a negative indirect effect on biomass (by reducing SLA through abundance shifts) and we could see some evidence for a negative direct effect (marginally significant, see Table S5). This means that sown SLA decreased aboveground biomass production, which was the opposite of our expectation (Lavorel & Grigulis, 2012). The effect is likely due to the better establishment of slow species at the beginning of the experiment. In the first year of the experiment, species suffered drought stress due to large amounts of bare ground before the species established themselves (Pichon pers obs). Slow growing, low SLA species may therefore have established better in the first year because of their higher investment in root biomass. This initial advantage for slow species means that plots with only fast species present (high sown SLA) still produce less biomass overall. However, the communities that were increasingly dominated by faster growing species, i.e. in which SLA increased due to abundance shifts, produced higher aboveground biomass. These contradictory results draw attention to the importance of initial establishment conditions (as in Mahaut et al., 2019 for instance) and suggest that relationships between effect traits and functions may change during community reassembly (Galland et al., 2019).

## Conclusion

Intraspecific changes in resource economics traits had large effects on overall functional composition, which were similar to abundance changes, showing that measuring traits *in situ* is important to accurately quantify functional composition. Intraspecific trait shifts tended to lead to convergence in functional composition between diverse communities, however, the degree of convergence depended on resource levels, diversity and enemy abundance and intra and interspecific shifts were opposing. Although large, intraspecific trait shifts did not translate into an effect on above or belowground biomass production. Intraspecific trait variation may therefore have different effects on function compared to interspecific trait differences, suggesting that intra and interspecific trait variation needs to be treated separately and not combined. Our results highlight the importance of a better understanding of how response and effect traits correlate at different scales and suggest that inter and intraspecific trait variation may have different consequences for ecosystem functioning.

## Acknowledgements

We would like to thank Hugo Vincent, Mervi Laitinen and Marlise Zimmermann, the technicians working with us on the PaNDiv Experiment, as well as the large team of helpers on the field and in the lab. This study was supported by funding of the Swiss National Science Foundation (31003A_160212).

## Authors contribution

N.A.P., S.L.C. and E.A. designed and set up the PaNDiv Experiment; N.A.P. and S.L.C. collected the data; N.A.P. analysed the data and wrote the first manuscript with substantial input from E.A and S.L.C. All authors contributed to revisions of the manuscript.

## Data accessibility

Once this manuscript is accepted, all the relevant data will be archived in figshare (https://figshare.com/).

