## Supplementary Information for "Intraspecific trait changes have large impacts on community functional composition but do not affect ecosystem function"

*bioRxiv*

Noémie A. Pichon, Seraina L. Cappelli, Eric Allan

**Table S1:** Subset of the structural equation model output for specific leaf area (SLA) and leaf dry matter content (LDMC). Non-conservative model, where the missing data for intraspecific shift are not replaced by monoculture values (see methods).

Effect of N enrichment, species richness (SR), fungicide (Fng), sown trait (Sown), abundance shift (Δ ab. shift) and intraspecific shift (Δ in. shift), on aboveground and belowground biomass production. N = 120 plots.

| Response | Predictor | Estimate | Std.Err | P(>\|z\|) | |
|---|---|---|---|---|---|
| Belowground biomass | N | 0.109 | 0.087 | 0.211 | |
| | SR | 0.030 | 0.086 | 0.728 | |
| | Fng | 0.058 | 0.087 | 0.503 | |
| | Δ in. shift SLA | 0.124 | 0.092 | 0.177 | |
| | Δ ab. shift SLA | -0.258 | 0.091 | 0.005 | ** |
| | Sown SLA | -0.248 | 0.092 | 0.007 | *** |
| Aboveground biomass | N | 0.303 | 0.086 | 0.000 | *** |
| | SR | -0.111 | 0.085 | 0.191 | |
| | Fng | 0.068 | 0.085 | 0.422 | |
| | Δ in. shift SLA | 0.097 | 0.091 | 0.286 | |
| | Δ ab. shift SLA | 0.108 | 0.090 | 0.227 | |
| | Sown SLA | -0.094 | 0.091 | 0.297 | |

| Response | Predictor | Estimate | Std.Err | P(>\|z\|) | |
|---|---|---|---|---|---|
| Belowground biomass | N | 0.118 | 0.085 | 0.164 | |
| | SR | 0.048 | 0.085 | 0.573 | |
| | Fng | 0.063 | 0.086 | 0.466 | |
| | Δ in. shift LDMC | -0.066 | 0.086 | 0.444 | |
| | Δ ab. shift LDMC | 0.142 | 0.088 | 0.108 | |
| | Sown LDMC | 0.359 | 0.087 | 0.000 | *** |
| Aboveground biomass | N | 0.309 | 0.084 | 0.000 | *** |
| | SR | -0.147 | 0.085 | 0.081 | . |
| | Fng | 0.050 | 0.085 | 0.559 | |
| | Δ in. shift LDMC | -0.124 | 0.085 | 0.147 | |
| | Δ ab. shift LDMC | -0.140 | 0.088 | 0.109 | |
| | Sown LDMC | -0.167 | 0.086 | 0.052 | . |

**Table S2:** For specific leaf area (SLA) and leaf dry matter content (LDMC), lmer outputs testing the presence of interactions between nitrogen (N), species richness (SR), fungicide (Fng) and sown traits on the difference between sown trait and intraspecific shift mean (Δ intraspecific shift), and sown trait and abundance shift mean. N = 120 plots.

| *Δ intraspecific shift SLA* | | | | |
|---|---|---|---|---|
| **Factor** | **Estimate** | **Std.Error** | **Pr(Chi)** | |
| (Intercept) | 0.004 | 0.258 | | |
| Nitrogen | 0.217 | 0.072 | | |
| Sown SLA | -0.186 | 0.088 | | |
| Species richness | 0.028 | 0.088 | | |
| Fungicide | -0.180 | 0.071 | | |
| Sown SLA x SR | 0.186 | 0.071 | 0.009 | ** |
| Sown SLA x SR | -0.191 | 0.088 | 0.028 | * |
| Sown SLA x Fng | -0.181 | 0.071 | 0.011 | * |

| *Δ abundance shift SLA* | | | | |
|---|---|---|---|---|
| **Factor** | **Estimate** | **Std.Error** | **Pr(Chi)** | |
| (Intercept) | 0.000 | 0.156 | | |
| Sown SLA | -0.359 | 0.156 | 0.021 | * |
| Species richness | -0.070 | 0.156 | | |
| Fungicide | 0.052 | 0.047 | | |
| SR x Fungicide | 0.098 | 0.047 | 0.039 | * |

| *Δ intraspecific shift LDMC* | | | | |
|---|---|---|---|---|
| **Factor** | **Estimate** | **Std.Error** | **Pr(Chi)** | |
| (Intercept) | 0.000 | 0.234 | | |
| Fungicide | 0.150 | 0.078 | 0.05 | . |

| *Δ abundance shift LDMC* | | | | |
|---|---|---|---|---|
| **Factor** | **Estimate** | **Std.Error** | **Pr(Chi)** | |
| (Intercept) | 0.000 | 0.131 | | |
| Nitrogen | -0.029 | 0.076 | | |
| Sown LDMC | -0.286 | 0.111 | 0.011 | * |
| Species richness | -0.096 | 0.111 | | |
| Fungicide | -0.159 | 0.076 | 0.034 | * |
| N x SR | 0.176 | 0.076 | 0.020 | * |

**Table S3:** Structural equation model output: effect of N enrichment, species richness (SR), fungicide (Fng) and sown specific leaf area (Sown SLA) on the shift of community mean SLA due to abundance shift (Δ ab. shift) and to intraspecific shift (Δ in. shift), and of sown SLA and the Δs on community weighted mean (CWM) SLA. N = 120 plots.

Model fit: Pvalue = 1.000, Chisq = 9.522, Df = 32, RMSEA = 0.000.

| Response | Predictor | Estimate | Std.Err | P(>\|z\|) | |
|---|---|---|---|---|---|
| CWM SLA | Sown SLA | 0.975 | 0.036 | 0.000 | *** |
| | Δ in. shift SLA | 0.436 | 0.034 | 0.000 | *** |
| | Δ ab. shift SLA | 0.538 | 0.035 | 0.000 | *** |
| Δ in. shift SLA | N | 0.144 | 0.082 | 0.077 | |
| | SR | 0.028 | 0.082 | 0.736 | |
| | Fng | -0.170 | 0.082 | 0.037 | |
| | Sown SLA | -0.186 | 0.082 | 0.023 | |
| | Sown SLA x N | 0.228 | 0.082 | 0.005 | ** |
| | Sown SLA x SR | -0.191 | 0.082 | 0.020 | * |
| | Sown SLA x Fng | -0.161 | 0.082 | 0.050 | * |
| Δ ab. shift SLA | N | 0.058 | 0.084 | 0.490 | |
| | SR | -0.070 | 0.084 | 0.408 | |
| | Fng | 0.052 | 0.084 | 0.535 | |
| | Sown SLA | -0.359 | 0.084 | 0.000 | *** |
| | SR x Fng | 0.097 | 0.084 | 0.249 | |

| Covariances | | Estimate | Std.Err | P(>\|z\|) |
|---|---|---|---|---|
| Sown SLA | Sown SLA x N | 0 | 0.09 | 1 |
| | Sown SLA x SR | 0.034 | 0.09 | 0.709 |
| | Sown SLA x Fng | 0 | 0.09 | 1 |
| N | Sown SLA x N | 0 | 0.09 | 1 |
| SR | Sown SLA x SR | -0.001 | 0.09 | 0.994 |
| Fng | Sown SLA x Fng | 0 | 0.09 | 1 |
| SR | Fng | 0 | | Fixed |
| N | SR | 0 | | Fixed |
| | Fng | 0 | | Fixed |
| Sown SLA | N | 0 | | Fixed |
| | Fng | 0 | | Fixed |
| SR x Fng | SR | 0 | 0.09 | 1 |
| | Fng | 0 | 0.09 | 1 |

| R square | |
|---|---|
| CWM SLA | 0.876 |
| Δ in. shift SLA | 0.201 |
| Δ ab. shift SLA | 0.150 |

**Table S4:** Structural equation model output: effect of N enrichment, species richness (SR), fungicide (Fng) and sown specific leaf area (Sown LDMC) on the shift of community mean LDMC due to abundance shift (Δ ab. shift) and to intraspecific shift (Δ in. shift), and of sown LDMC and the Δs on community weighted mean (CWM) LDMC. N = 120 plots.

Model fit: Pvalue = 0.999, Chisq = 14.865, Df = 35, RMSEA = 0.000.

| Response | Predictor | Estimate | Std.Err | P(>\|z\|) | |
|---|---|---|---|---|---|
| CWM LDMC | Sown LDMC | 0.599 | 0.048 | 0.000 | *** |
| | Δ in. shift LDMC | 0.528 | 0.046 | 0.000 | *** |
| | Δ ab. shift LDMC | 0.626 | 0.048 | 0.000 | *** |
| Δ in. shift LDMC | N | -0.045 | 0.090 | 0.616 | |
| | SR | -0.070 | 0.090 | 0.439 | |
| | Fng | 0.124 | 0.090 | 0.168 | |
| | Sown LDMC | -0.080 | 0.090 | 0.376 | |
| Δ ab. shift LDMC | N | -0.016 | 0.084 | 0.851 | |
| | SR | -0.096 | 0.084 | 0.250 | |
| | Fng | -0.168 | 0.084 | 0.045 | * |
| | Sown LDMC | -0.286 | 0.084 | 0.001 | ** |
| | N x SR | 0.184 | 0.084 | 0.029 | * |

| Covariances | | Estimate | Std.Err | P(>\|z\|) |
|---|---|---|---|---|
| Sown LDMC | Sown LDMC x N | 0 | 0.088 | 1 |
| | Sown LDMC x SR | -0.231 | 0.092 | 0.013 |
| | Sown LDMC x Fng | 0 | 0.088 | 1 |
| N | Sown LDMC x N | 0 | 0.09 | 1 |
| SR | Sown LDMC x SR | 0.011 | 0.088 | 0.9 |
| Fng | Sown LDMC x Fng | 0.000 | 0.090 | 1 |
| SR | Fng | 0 | | Fixed |
| N | SR | 0 | | Fixed |
| | Fng | 0 | | Fixed |
| Sown LDMC | N | 0 | | Fixed |
| | Fng | 0 | | Fixed |
| N x SR | SR | 0 | 0.09 | 1 |
| | N | 0 | 0.09 | 1 |

| R square | |
|---|---|
| CWM LDMC | 0.754 |
| Δ in. shift LDMC | 0.029 |
| Δ ab. shift LDMC | 0.154 |

**Table S5:** Structural equation model output: effect of N enrichment, species richness (SR), fungicide (Fng) and sown specific leaf area (Sown SLA) on the shift of community mean SLA due to abundance shift (Δ ab. shift) and to intraspecific shift (Δ in. shift), and of sown SLA, N enrichment, SR, Fng and the Δs on aboveground and belowground biomass production. N = 120 plots.

Model fit: Pvalue = 1.000, Chisq = 7.7733, Df = 33, RMSEA = 0.000.

| Response | Predictor | Estimate | Std.Err | P(>\|z\|) | |
|---|---|---|---|---|---|
| Belowground biomass | N | 0.112 | 0.087 | 0.197 | |
| | SR | 0.029 | 0.085 | 0.734 | |
| | Fng | 0.054 | 0.086 | 0.534 | |
| | Δ in. shift SLA | 0.120 | 0.091 | 0.187 | |
| | Δ ab. shift SLA | -0.295 | 0.092 | 0.001 | ** |
| | Sown SLA | -0.249 | 0.092 | 0.007 | ** |
| Aboveground biomass | N | 0.295 | 0.083 | 0.000 | *** |
| | SR | -0.107 | 0.082 | 0.192 | |
| | Fng | 0.065 | 0.083 | 0.430 | |
| | Δ in. shift SLA | 0.088 | 0.087 | 0.315 | |
| | Δ ab. shift SLA | 0.192 | 0.088 | 0.029 | * |
| | Sown SLA | -0.146 | 0.089 | 0.098 | . |
| Δ in. shift SLA | N | 0.163 | 0.080 | 0.042 | |
| | SR | 0.008 | 0.080 | 0.916 | |
| | Fng | -0.151 | 0.080 | 0.060 | |
| | Sown SLA | -0.167 | 0.080 | 0.037 | |
| | Sown SLA x N | 0.209 | 0.081 | 0.009 | ** |
| | Sown SLA x SR | -0.173 | 0.081 | 0.032 | * |
| | Sown SLA x Fng | -0.180 | 0.081 | 0.026 | * |
| Δ ab. shift SLA | N | 0.065 | 0.085 | 0.444 | |
| | SR | -0.076 | 0.085 | 0.367 | |
| | Fng | 0.059 | 0.085 | 0.488 | |
| | Sown SLA | -0.353 | 0.085 | 0.000 | *** |
| | SR x Fng | 0.104 | 0.085 | 0.221 | |

| Covariances | | Estimate | Std.Err | P(>\|z\|) |
|---|---|---|---|---|
| Sown SLA | Sown SLA x N | 0.008 | 0.09 | 0.927 |
| | Sown SLA x SR | 0.025 | 0.091 | 0.782 |
| | Sown SLA x Fng | 0.008 | 0.09 | 0.927 |
| N | Sown SLA x N | 0.008 | 0.091 | 0.926 |
| SR | Sown SLA x SR | 0.008 | 0.09 | 0.931 |
| Fng | Sown SLA x Fng | 0.008 | 0.091 | 0.926 |
| N | Fng | 0 | | Fixed |
| | Sown SLA | 0 | | Fixed |
| Sown SLA | Fng | 0 | | Fixed |
| Belowground biomass | Aboveground biomass | -0.118 | 0.076 | 0.12 |

| R square | |
|---|---|
| Belowground biomass | 0.132 |
| Aboveground biomass | 0.212 |
| Δ in. shift SLA | 0.194 |
| Δ ab. shift SLA | 0.148 |

**Table S6:** Structural equation model output: effect of N enrichment, species richness (SR), fungicide (Fng) and sown specific leaf area (Sown LDMC) on the shift of community mean LDMC due to abundance shift (Δ ab. shift) and to intraspecific shift (Δ in. shift), and of sown LDMC, N enrichment, SR, Fng and the Δs on aboveground and belowground biomass production. N = 120 plots.

Model fit: Pvalue = 1.000, Chisq = 1.479, Df = 12, RMSEA = 0.000.

| Response | Predictor | Estimate | Std.Err | P(>\|z\|) | |
|---|---|---|---|---|---|
| Belowground biomass | N | 0.112 | 0.085 | 0.188 | |
| | SR | 0.051 | 0.085 | 0.550 | |
| | Fng | 0.061 | 0.087 | 0.483 | |
| | Δ in. shift LDMC | -0.066 | 0.087 | 0.445 | |
| | Δ ab. shift LDMC | 0.208 | 0.090 | 0.021 | * |
| | Sown LDMC | 0.351 | 0.089 | 0.000 | *** |
| Aboveground biomass | N | 0.316 | 0.084 | 0.000 | *** |
| | SR | -0.146 | 0.085 | 0.086 | . |
| | Fng | 0.051 | 0.086 | 0.557 | |
| | Δ in. shift LDMC | -0.103 | 0.086 | 0.233 | |
| | Δ ab. shift LDMC | -0.157 | 0.090 | 0.080 | . |
| | Sown LDMC | -0.107 | 0.088 | 0.224 | |
| Δ in. shift LDMC | N | -0.059 | 0.090 | 0.513 | |
| | SR | -0.057 | 0.090 | 0.529 | |
| | Fng | 0.110 | 0.090 | 0.218 | |
| | Sown LDMC | -0.070 | 0.090 | 0.435 | |
| Δ ab. shift LDMC | N | -0.014 | 0.085 | 0.864 | |
| | SR | -0.098 | 0.085 | 0.248 | |
| | Fng | -0.167 | 0.085 | 0.048 | * |
| | Sown LDMC | -0.287 | 0.084 | 0.001 | *** |
| | N x SR | 0.186 | 0.085 | 0.029 | * |

| Covariances | | Estimate | Std.Err | P(>\|z\|) |
|---|---|---|---|---|
| N | Fng | 0 | | Fixed |
| Sown LDMC | SR | 0.041 | 0.091 | 0.65 |
| Belowground biomass | Aboveground biomass | -0.081 | 0.078 | 0.295 |
| N x SR | SR | 0.008 | 0.091 | 0.926 |
| | N | -0.008 | 0.091 | 0.926 |

| R square | |
|---|---|
| Belowground biomass | 0.145 |
| Aboveground biomass | 0.159 |
| Δ in. shift LDMC | 0.024 |
| Δ ab. shift LDMC | 0.153 |

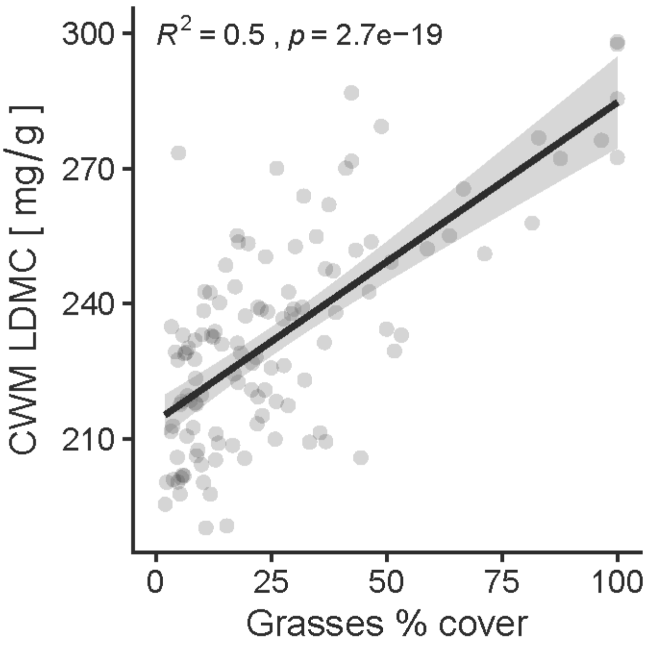


**Figure S1:** Sown grasses relative percentage cover compared to sown herbs in August 2017. Pearson correlation with the community weighted mean leaf dry matter content (CWM LDMC) value per plot. Correlation coefficient 0.71, $R^2$ 0.5, pvalue <0.001. N = 200 plots.
